# Transcriptomic, Genomic, and Clinical Characterization of Morphological Classes in Localized and Metastatic Pancreatic Cancer

**DOI:** 10.64898/2026.04.07.717011

**Authors:** Eugenia Flores Figueroa, Yuanchang Fang, Ayah Elqaderi, Maryam Monajemzadeh, Amy Zhang, Gun Ho Jang, Michelle Chan-Seng-Yue, Karen Ng, Tom Ouellette, Stephanie Ramotar, Daniela Bevacqua, Shawn Hutchinson, Yuqiong Rachel Ding, Sheng-Beng Liang, Syeda Mariam Hasnain, Grainne M. O’Kane, Sandra Fischer, Klaudia Nowak, Barbara Grünwald, Anna Dodd, Julie M Wilson, Erica S Tsang, Steven Gallinger, Jennifer J. Knox, Faiyaz Notta, Robert C Grant

## Abstract

**Background:** Histomorphology is a strong prognostic biomarker correlated with basal-like and classical programs in surgically resected pancreatic ductal adenocarcinoma (PDAC). However, the spectrum of morphology and its biological associations remain poorly defined in advanced disease.

**Objectives:** We explored the transcriptomic and genomic underpinnings and clinical relevance of morphological classes across localized and metastatic PDAC.

**Design:** We unified morphological classifications into four classes: glandular, cribriform, solid, and squamous. We integrated transcriptome and whole-genome sequencing following laser-capture microdissection with morphological classifications in 348 PDAC patients, where half of the cohort included locally advance and metastatic stages to uncover molecular associations.

**Results:** Non-glandular morphologies comprised three distinct classes that were enriched in metastatic disease. Transcriptomic profiling exhibited that glandular tumours predominantly expressed classical epithelial programs, although a subset displayed partial or full epithelial– mesenchymal transition signatures. In contrast, non-glandular morphologies showed basal-like transcriptional programs with subtype-specific pathways, including ciliogenesis in cribriform tumours, extracellular matrix remodelling and immune evasion in solid tumours, and keratinisation programs in squamous tumours. The solid class was significantly enriched in liver metastatic lesions and was associated with increased intra-tumoural morphological heterogeneity, whole-genome doubling, *KRAS* major allelic imbalance, and elevated KRAS–ERK signalling.

**Conclusion:** Non-glandular morphologies identify biologically distinct PDAC tumour states that are enriched in liver metastases and associated with subtype-specific transcriptional programs and KRAS-driven genomic alterations.

## Introduction

Pancreatic ductal adenocarcinoma (PDAC) is a deadly disease with a rising incidence^1^. Despite intensive study, the fundamental causes of the aggressive and treatment-resistant nature of PDAC remain unknown. Histopathological morphology of PDAC has long been recognized as a strong prognostic biomarker. Recent studies demonstrated that morphological differences in primary PDAC reflect intrinsic biology. For example, primary tumours with well-differentiated, glandular morphologies are associated with classical gene expression programs, while poorly differentiated non-glandular and squamous phenotypes are associated with basal-like programs^2–4^. However, glandular *versus* non-glandular classification encompasses only one dimension of tumour morphology, and emerging evidence suggests that basal-like programs exist along a continuum, heavily influenced by cellular plasticity and epithelial-mesenchymal transition (EMT) processes^5,6^. Most importantly, molecular studies on PDAC morphology to date have used small cohorts and focused mostly on early-stage primary tumours, despite 80% of PDAC presenting at an advanced stage^2–4,7^. Late-stage cancers have distinct biology, for example, more frequent basal-like programs and whole-genome doubling^8^. However, comprehensive characterization of the molecular mechanisms underlying PDAC morphology requires a large cohort spanning the spectrum of disease.

In this study, we present the largest cohort to date integrating histological morphology with matched transcriptomic, genomic, and clinical data across surgical and unresectable advanced PDAC. We unified published morphological classifications and described intra- and inter-patient variation. We then integrated transcriptomic and genomic data from a tumour-enriched cohort using laser microdissection (LCM) to delineate the molecular basis of the morphological classes.

## Results

### Morphological Classes in Pancreatic Cancer

We compiled previously published tumour histology categories from resected cases^2,4,7^ and extended this classification to metastatic disease. These were unified into four morphological classes: glandular, cribriform, solid, and squamous (see Methods, **Figure 1A-B, Supplementary Figure 1**), which span a continuum of gland-forming to non-gland forming architectures. For each case, morphologic patterns were semi-quantitatively estimated as a percentage of total tumour area, and the dominant pattern was assigned as the primary morphologic class, except for squamous morphology (see Methods). To determine whether these morphological classes are retained in metastatic disease, we assembled a cohort with matched histological slides, whole transcriptome and whole genome sequencing data (n = 348), comprising 38% with a resected tumour (Stage I and II), 7% with a locally advanced disease (Stage III), 46% with distant metastases (Stage IV), and 9% with missing staging information. We additionally analyzed 8 tissue microarrays (TMA) containing tumours from 181 donors across all clinical stages (**Figure 1A**). The glandular class included all glandular-forming morphologies from previous descriptions^2,4,7^. The glandular class is characterized by well-defined tubular structures of variable sizes with a patent lumen with or without papillary projection. These structures are lined by pancreaticobiliary-type or foveolar-gastric epithelium, with eosinophilic to amphophilic cytoplasm and low to moderate cytological atypia. The non-glandular class represents the progressive loss of glandular differentiation and the acquisition of mesenchymal or squamoid features. The cribriform subtype, as its name implies, describes the well-known cribriform appearance in histopathology, which is a group of cohesive sheets or nests of tumour cells showing round punched out spaces, without true epithelial lining (pseudo-lumen). This class also includes signet ring morphology, and poorly formed and angulated glands. Tumour cells in this subtype retain some glandular cytomorphology (the cells still of a pancreatobiliary or gastric type) but no well-formed glands with tubular lumens are identified (**Figure 1B, Supplementary Figure 1**). The solid subtype described tumour cells arranged in sheets, nests, cords, and isolated cells with moderate to severe atypia. These tumour cells completely lost their glandular differentiation cytologically and architecturally, lacking both mucin and true lumen. The tumour cells in the squamous subtype are defined by the presence of squamoid differentiation, which is characterized mainly by the presence of polygonal cells, with abundant eosinophilic cytoplasm, and intercellular bridges (**Figure 1B, Supplementary Figure 1**). The squamous differentiation can be moderate to poorly differentiated, where the cells can show minimal to no keratinization (**Figure 1B, Supplementary Figure 1**). For each tumour, the dominant morphology was determined as the one with the maximum percentage among glandular, cribriform and solid subtypes. For the squamous subtype we used a cut off of >30%. Downstream analysis used this dominant morphology, except for the analysis of tumour heterogeneity.

**Figure 1.**
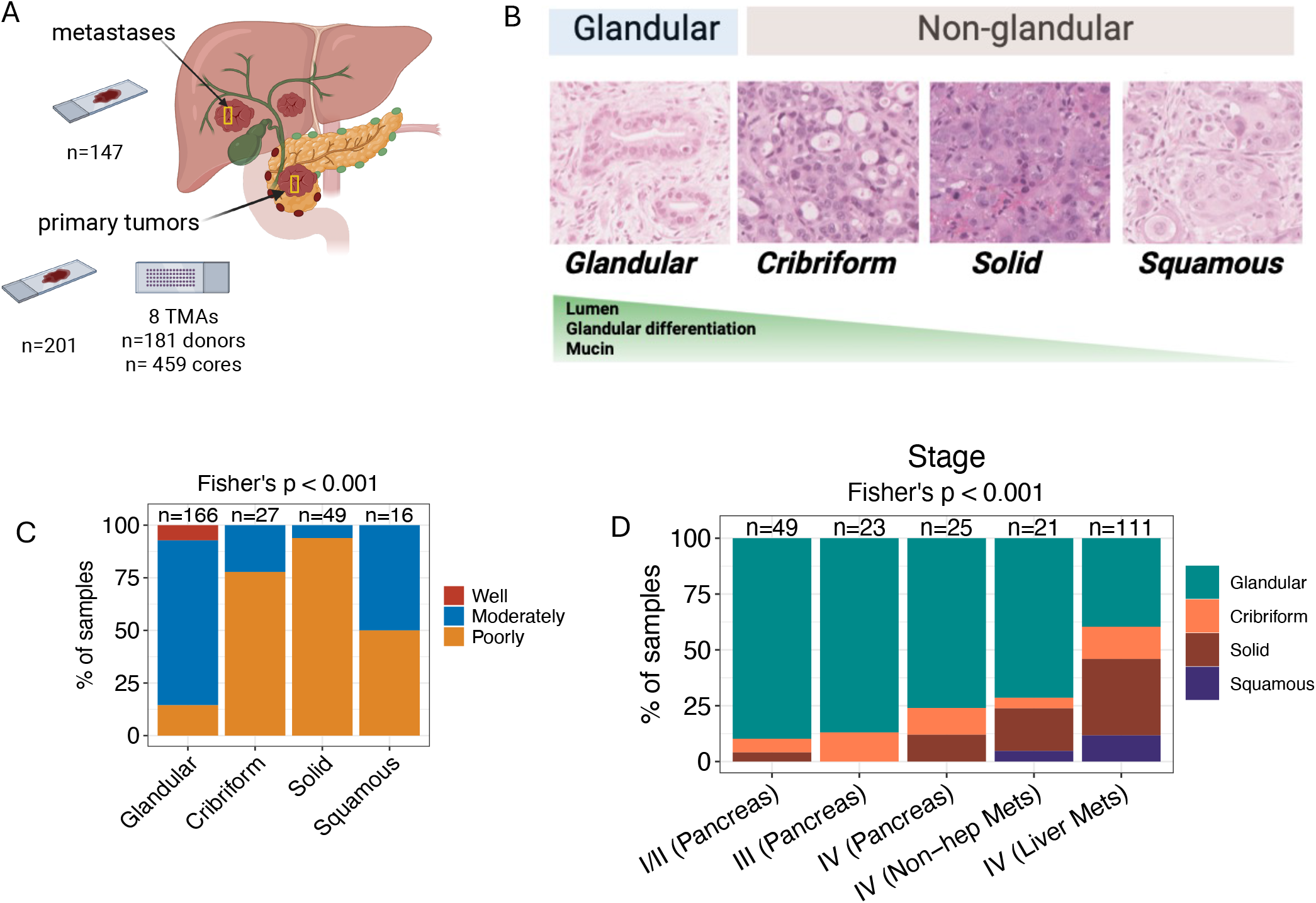
Profiling morphological subtypes in resectable and advanced pancreatic cancer. (A) Sample composition of the study cohort. Metastatic samples were obtained from the liver, lung, peritoneum, and other sites. TMA, tissue micro-array. (B) Representative images illustrating the morphological classification used in this study. Hematoxylin and Eosin (H&E) stain liver biopsies infiltrated by metastatic pancreatic tumours illustrating distinct morphological subtypes, including glandular and non-glandular (cribriform, solid, and squamous) morphologies. (C) Stacked bar plots showing the distribution of morphological subtypes according to World Health Organization (WHO) differentiation grade and (D) clinical stage with tissue site.

As expected, morphologies were strongly associated with reported tumour differentiation (n = 258; p <0.001), evaluated according to the World Health Organization (WHO) criteria^9^, which defines well differentiated tumours as displaying more than 90% glandular morphology, moderately differentiated tumours as displaying 10-90% glandular morphology and poorly differentiated tumours as displaying with less than 10% glandular morphology. Most (78%) of glandular tumours were moderately differentiated, while most (50%-94%) of the non-glandular tumours were poorly differentiated (**Figure 1C**).

PDAC is characterized by a dense desmoplastic tumour microenvironment, so we next investigated if tumour cellularity differs across morphologies. Primary pancreatic tumours had significantly lower cellularity, compared to metastases (30% vs. 73%) (Supplementary **Figure 2A**). Tumour cellularity was not associated with morphologies in primary tumours (p = 0.34), while in metastases, tumour cellularity was in general higher in non-glandular tumours than the glandular ones (p = 0.02) (Supplementary **Figure 2A**).

### Liver Metastases are Enriched in Non-Glandular Morphological Classes

The distribution of morphological classes varied significantly by stage (p < 0.001), with glandular morphology predominating in stage I/II tumours (>80%) but comprising only ∼50% of stage III/IV disease (**Supplementary Figure 2B**). When metastatic cases were further stratified by site, enrichment of non-glandular morphologies was observed predominantly in liver metastases, whereas non-hepatic metastases exhibited a distribution more similar to pancreatic tumours from the same stage (IV) (**Figure 1D**). This difference was largely driven by an increased incidence of solid morphology in liver metastases. These findings should be interpreted in the context of the smaller non-hepatic metastatic cohort (n = 21) relative to liver metastases (n = 111). Notably, stage III tumours were almost exclusively glandular, with no solid or squamous morphologies observed. No differences in the distribution of morphological subtypes were detected with respect to sex or race (**Supplementary Figure 2C–D**).

### Morphological Heterogeneity Increases with Metastatic Spread

Metastatic tumours displayed greater morphological heterogeneity than primary tumours. To quantify this, we analyzed tumour heterogeneity within a single histological section (intra-tumour) from primary pancreatic tumours (n = 115) and metastatic lesions (n = 143) which predominately originated in the liver (123/143) (**Figure 2A**). We found that 82% of primary pancreatic tumours exhibited a single morphologic class (defined as pure), whereas metastatic tumours were more heterogeneous, with 52% displaying multiple morphological classes (defined as mixed) (**Figure 2B**). Across primary and metastatic sites, glandular tumours were the most homogeneous, with 68% classified as pure. In contrast, cribriform tumours were the most heterogeneous, with only 9% classified as pure. Solid and squamous tumours showed a balanced distribution between pure and mixed classes (**Figure 2C**). We next compared the proportion and co-occurrence of each morphological class across different stages (I-IV) (**Figure 2D**). Most mixed cases had a combination of cribriform-glandular or cribriform-solid morphologies. Squamous cases were mainly pure or mixed with glandular morphologies. Most stage I/II cases were pure glandular, while stage IV was mixed or pure non-glandular. Collectively, these analyses reveal a morphologic continuum across disease stages, from predominantly pure glandular tumours in stage I/II to increasingly heterogeneous, non-glandular compositions in stage IV.

**Figure 2.**
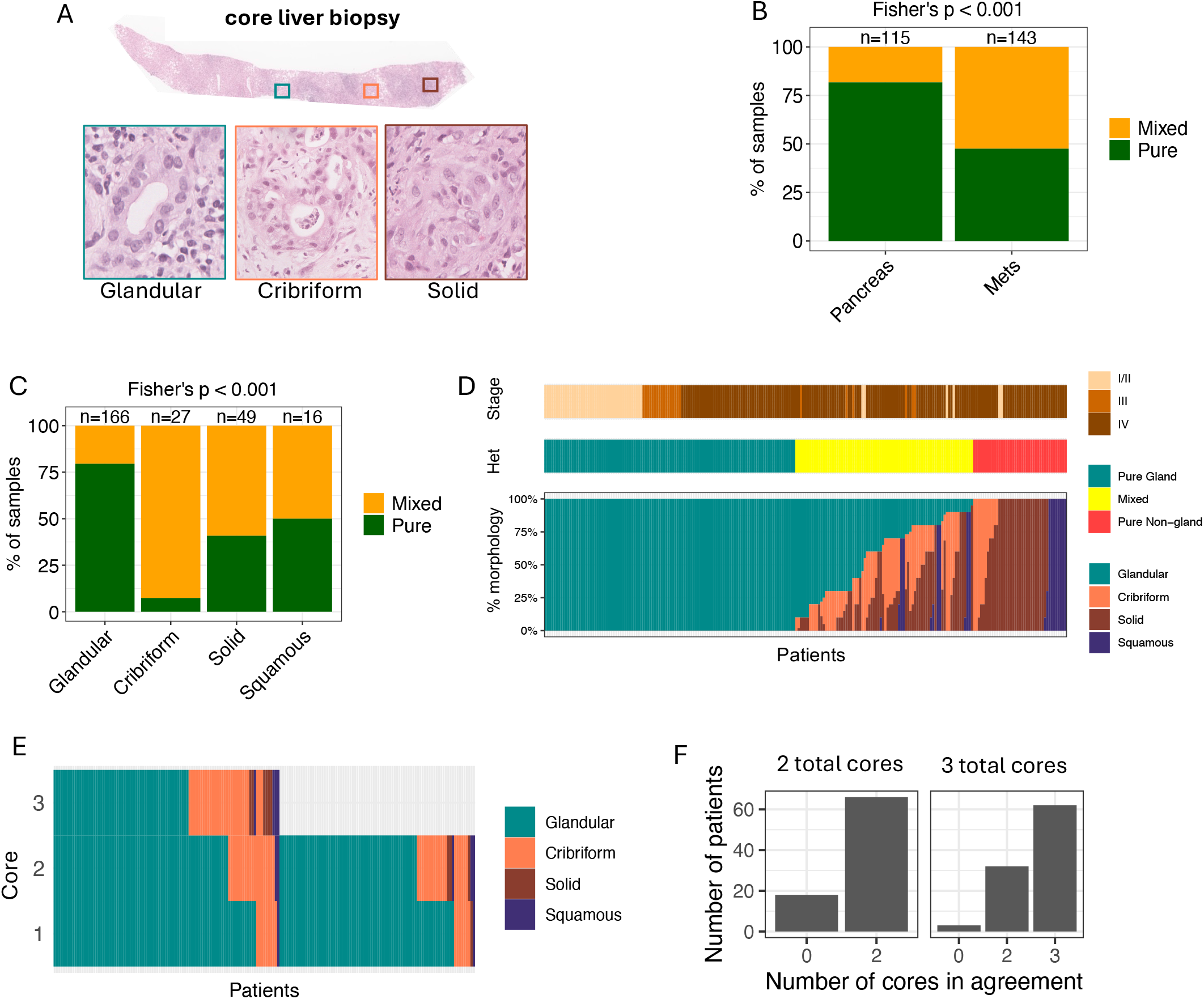
Histomorphological heterogeneity in pancreatic ductal adenocarcinoma (PDAC). (A) Representative Hematoxylin and Eosin (H&E) stained slide from liver biopsies infiltrated by metastatic pancreatic tumours illustrating three distinct morphological subtypes. Stacked bar plots show the proportion of pure (single morphology) and mixed (two or more morphologies) tumours according to (B) tumour site (pancreas or metastasis (Mets)) and (C) morphological subtypes. (D) Stacked bar plot depicting the percentage of each morphological subtype per patient, with tracks indicating clinical stage and heterogeneity status. (E) Stacked bar plot summarising 8 tissue microarrays with 459 cores from 191 patients. Only patients with 2-3 cores were included in the analysis, and the dominated morphology was determined per core. (F) Number of cores per patient exhibiting concordant morphological classification.

To expand our observations, we use 8 tissue microarrays (TMAs) comprising 459 cores from 181 resectable patients from all clinical cases but predominantly resectable. Each case was represented by at least two or three independent cores (**Figure 2A**). All four morphological classes were present, and similar patterns on morphological distribution were observed. Most (77%) cores from TMAs displayed glandular morphology (**Figure 2E**). Despite inter-tumour-core heterogeneity, we observed a high concordance (88% with at least 2 cores in agreement) among cores from the same patient (**Figure 2F**).

### Non-Glandular Tumours Display Distinctive Basal-like Transcriptional Programs, while a Subset of Glandular Tumours Display EMT Programs

To define the molecular features of the morphological classes, we conducted differential gene expression analysis using 230 paired tumour RNA-seq (**Supplementary Table 2**), followed by overrepresentation analysis of pathways (**Supplementary Table 3**, Methods). Glandular tumours were enriched in genes related to classical signatures, epithelial identity and cell adhesion, such as *LGALS4, CEACAM6, CLDN18* and *TFF1* (**Supplementary Figure 3A**). Pathway analysis confirmed the epithelial nature of glandular tumours with overrepresented signatures related to gastric cell fate (**Figure 3A**). Overall, glandular tumours expressed higher classical (**Figure 3B, C**) and lowest basal (**Figure 3B, D**) signatures, in agreement with earlier publications using resected cohorts^2,4^. Interestingly, a subset of glandular tumours exhibited high expression of (partial) epithelial-mesenchymal transition (EMT)^10^ (**Figure 3B, E**) and full EMT or mesenchymal^10,11^ (**Figure 3B, F**) gene signatures. To investigate whether the heterogeneity in the expression of EMT signatures in early stages was driven by differences in tumour cellularity between resected and metastatic samples, we compared EMT signatures from early and advanced tumours within the pancreas. We found that early-stage (I/II) glandular and non-glandular tumours expressed higher levels of EMT hallmark signature (**Figure 3G**) than late-stage tumours (III and IV). These results demonstrate that there is a subset of samples in early-stage pancreatic cancer that harbours high EMT scores even in well differentiated gland-forming tumours. These observations underscore the intrinsically heterogeneous nature of glandular tumours and suggest an early activation of partial EMT and EMT programs.

**Figure 3.**
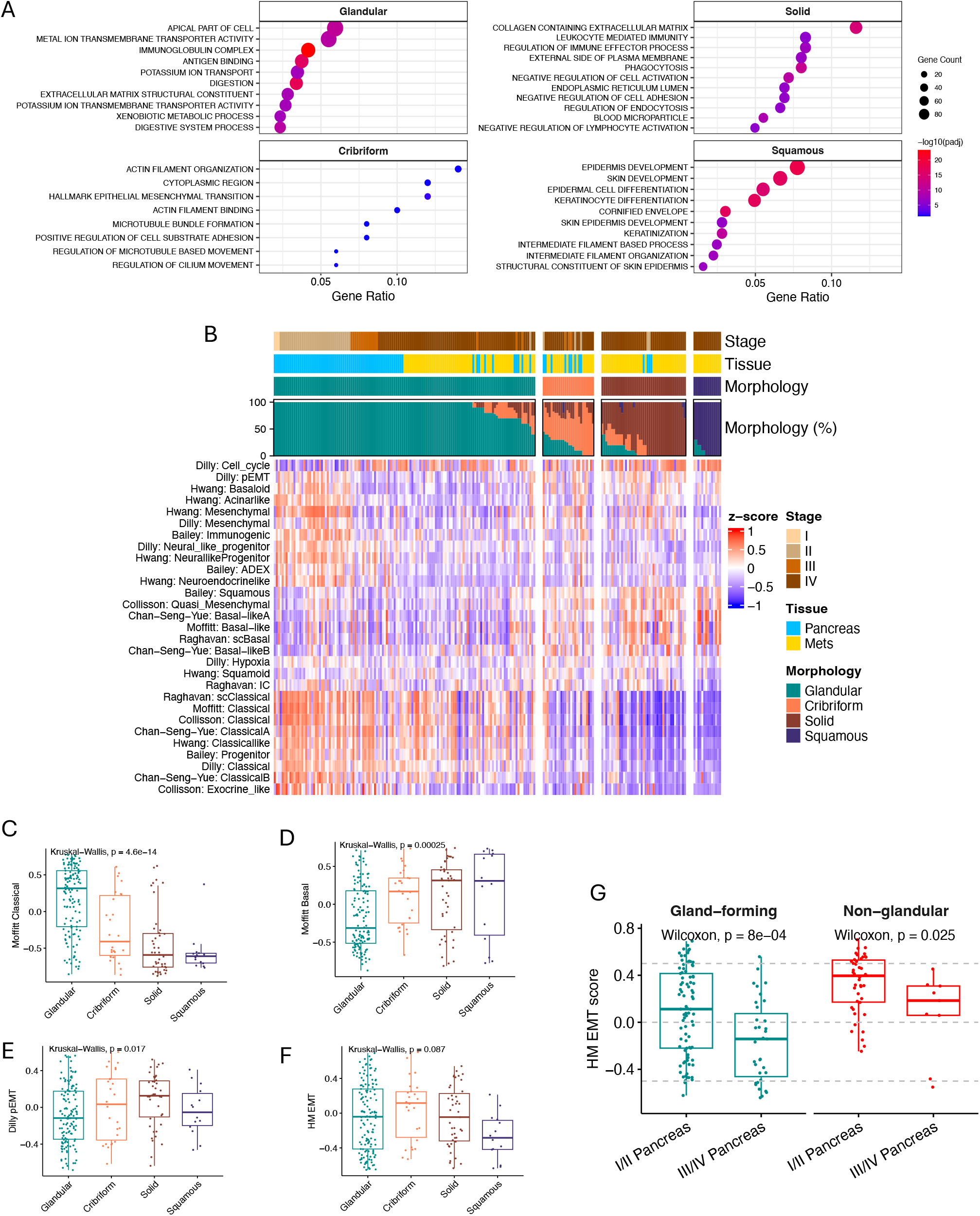
Transcriptomic characterization of morphological subtypes in pancreatic adenocarcinoma (PDAC). (A) Overrepresented gene sets identified from marker genes associated with each morphological subtype. (B) Scores of previously published transcriptional subtypes in PDAC calculated using Gene Set Variation Analysis (GSVA). (C-F) Bar plots comparing GSVA scores of selected gene sets from (C, D) Moffitt et al.^24^, (E) Dilly et al.^10^ and (F) Hallmark Epithelial-Mesenchymal Transition (EMT)^25^ in the tumour from all stages combined. (G) Bar plots showing GSVA scores of Hallmark EMT in primary tumours only according to stage. ^8^

Non-glandular morphological classes have distinct programs. Although non-glandular morphologies expressed similar levels of basal signatures (**Figure 3B, D**), they had differential expression of cell cycle marker genes (**Supplementary Figure 3B**) and biological pathways like keratinization (**Supplementary Figure 3C)**, neural-like progenitor **(Supplementary Figure 3D)**, among others (**Figure 3B**), including low levels or lack of classical signatures (**Figure 3C**). Cribriform tumours were characterized by genes associated with differentiation (*TAGLN, TPPP3*, and *VGLL3*), metabolism (*SLC16A5* and *PVT1*), and interactions with the tumour microenvironment (*PECAM1, CCL21*, and *ANGPTL2*) (**Table 2, Supplementary Figure 3A**), which converge on pathways of EMT and ciliogenesis (**Figure 3A**). The solid morphology exhibited pathways for extracellular matrix remodeling, nutrient uptake, and immune evasion, associated with genes including *CALB1, ITGB3, TIMP3, ZEB2* and *SERPINE1* (**Supplementary Figure 3A**), and hallmark pathways associated with extracellular matrix and cell adhesion, which can facilitate metastasis and immune regulation, reflecting adaptations to metastatic environments (**Figure 3A**). The squamous morphology was linked to genes involved in keratinization such as *KRT15, FAT2, NECTIN1, KRT6* and *TEAD2*, and developmental signaling that enhanced growth and survival (**Supplementary Figure 3A**). Principal component analysis (PCA) on the marker genes demonstrated a transcriptomic continuum from glandular, cribriform, solid and finally to squamous morphology (**Supplementary Figure 3E**). Taken together, these results demonstrate a morphological and transcriptional continuum underlying pancreatic cancer progression, rather than isolated, fixed categories. Pancreatic cancer may evolve along a spectrum where cribriform morphology sits between glandular, well-differentiated and non-glandular classes, both histologically and transcriptionally.

### Whole-Genome Doubling is Frequent in Non-Glandular Tumours

We next examined whether genomic alterations differed across morphological subtypes. Whole-genome doubling is a common event in PDAC and has been associated with more aggressive behavior^12–15^. In advanced PDAC, non-glandular tumours had a higher frequency of polyploid genomes (p = 0.03). Solid morphology showed the higher increased in frequency in the polyploid in comparison to the diploid group (**Figure 4B**), emphasizing the association between increased ploidy and poorly differentiated morphologies. In resectable PDAC, non-glandular cases were too small for meaningful comparison (p = 0.79) (**Figure 4A**). No significant differences in tumour mutation burden (TMB) were observed between morphological subtypes in either resectable (p = 0.23) or advanced tumours (p = 0.63) (**Figure 4C**). Across all stages, homologous recombination deficiency (HRD) was not associated with morphologies, but cribriform tumours trended towards a higher HRD frequency in advanced cases and solid classes to non-HRD, which merits further investigation in a larger cohort (p = 0.056) (**Figure 4D,E**). Next, we examined whether alterations in canonical cancer driver genes, including mutations, loss of heterozygosity, amplifications, and deletions, differed across morphological subtypes (**Figure 4F**). Among these, *MYC* showed a significant association with morphology, with higher amplification (copy number > 3 x ploidy) frequency in non-glandular compared with glandular tumours (15% vs. 5%, Fisher’s exact test, p = 0.015) (**Figure 4F**).

**Figure 4.**
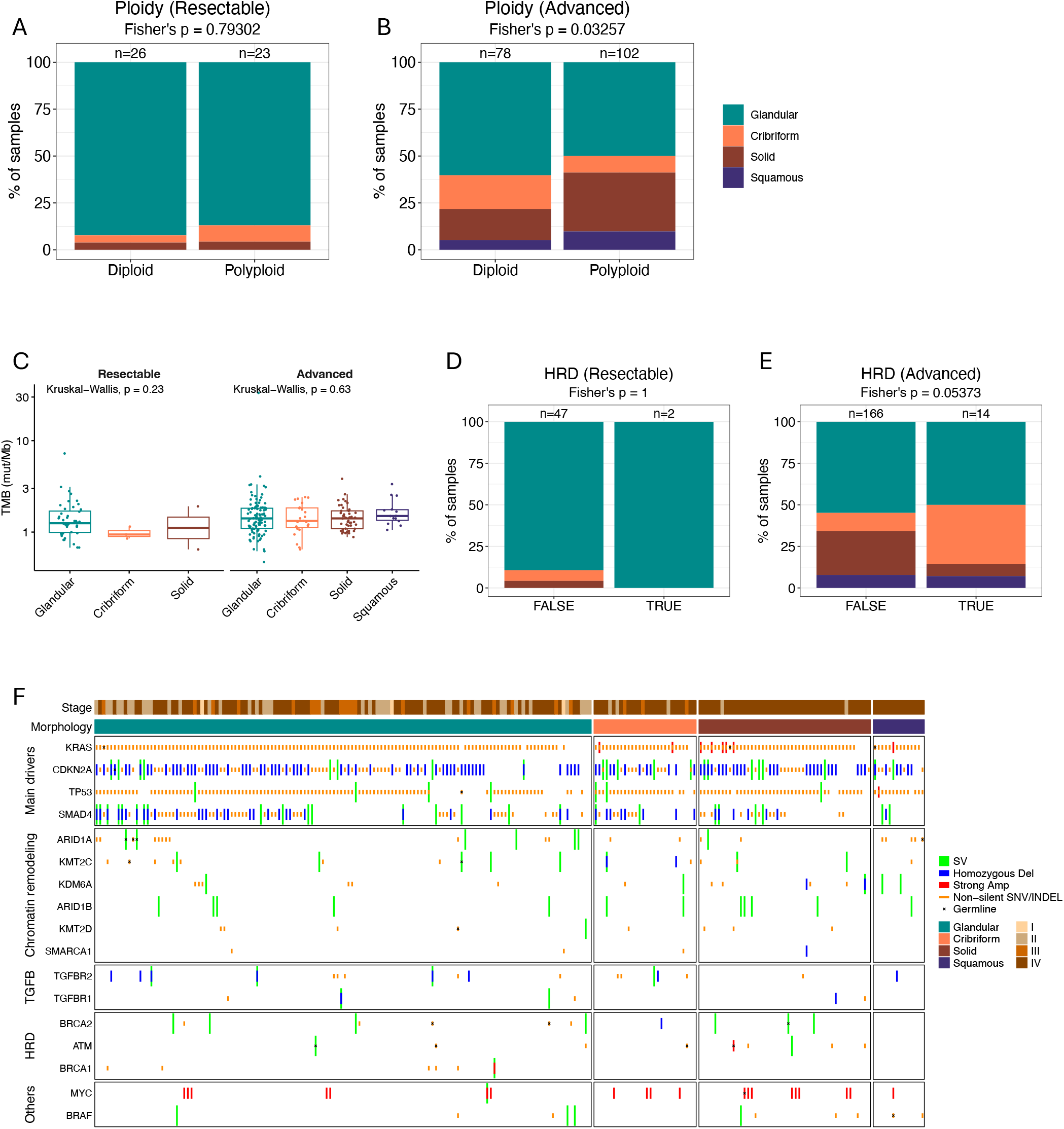
Genomic landscape of morphological subtypes in pancreatic ductal adenocarcinoma (PDAC). Correlations between tumour ploidy and morphology in (A) resectable and (B) advanced disease. (C) Tumour mutation burden (TMB), calculated as the number of nonsynonymous single-nucleotide variants (SNVs) and frame-shift small insertions and deletions (INDELs) within the coding regions of the human genome (∼34 Mb). Association between tumour homologous recombination deficiency (HRD) status and morphology in (D) resectable and (E) advanced disease. (F) Oncoprint depicting alterations in key driver genes. SV, structural variation; Del, deletion; Strong AMP (amplification), amplification defined as gene copy number greater than three times tumour ploidy.

### Non-glandular Morphologies Harbour KRAS Allelic Imbalance and Increased KRAS-ERK Signaling

*KRAS* is the most frequently mutated driver gene in PDAC, with specific alleles and allelic imbalance playing critical roles in tumour initiation, prognosis, and response to KRAS inhibitors^10,16–19^. We explored the contribution of *KRAS* to morphological progression by analyzing whole-genome sequencing. We focus the analysis on advanced cases, as non-glandular morphologies are rare in resectable cases. First, we assessed *KRAS* imbalance^13^, given its strong prognostic associations^20^. In *KRAS*-mutant tumours, non-glandular morphologies, particularly solid morphology, strongly correlate with major *KRAS* imbalances (p = 0.001) (**Figure 5A**). Aligned with this, we observed that *KRAS* copy number increased along the morphological spectrum from glandular to squamous tumours (p = 0.00032) **(Figure 5B**). To assess the transcriptional activation of the KRAS pathway, we used a KRAS-ERK signaling gene set specifically developed for PDAC^21^. Accordingly, elevating KRAS-ERK singling was found along the morphological spectrum (p = 0.00016) **(Figure 5C)**. These data suggest that in *KRAS* mutant tumours, the gains in *KRAS* genomic dosage may contribute to morphological progression towards more undifferentiated non-glandular tumours via activating the KRAS-ERK signaling. Notably, in the *KRAS* wild-type cases, there were more non-glandular tumours than glandular tumours (7 vs. 4) (**Figure 5A**). We found that the *KRAS* wild-type tumours lacked strong *KRAS* amplifications (total copies > 4) (**Figure 5E**) but sustained a similar level of KRAS-ERK signaling as the *KRAS* mutant tumours did (**Figure 5F**). These imply that *KRAS* wild-type tumours may still drive morphological progression by sustaining downstream ERK signaling, while this process was not achieved by increasing *KRAS* genomic dosage, but through other mechanisms independent of mutant *KRAS*, which warrants further investigation. Finally, we analyzed *KRAS* alleles across the morphological subtypes. We did not observe any significant association between a specific *KRAS* allele and a morphological subtype, but we identified two trends: *KRAS* p.G12R was enriched in glandular tumours, while *KRAS* p.Q61H was enriched in non-glandular morphologies, including squamous (**Figure 4G**). Together, these findings link KRAS dosage and downstream ERK signaling to the emergence of non-glandular morphologies in metastatic PDAC.

**Figure 5.**
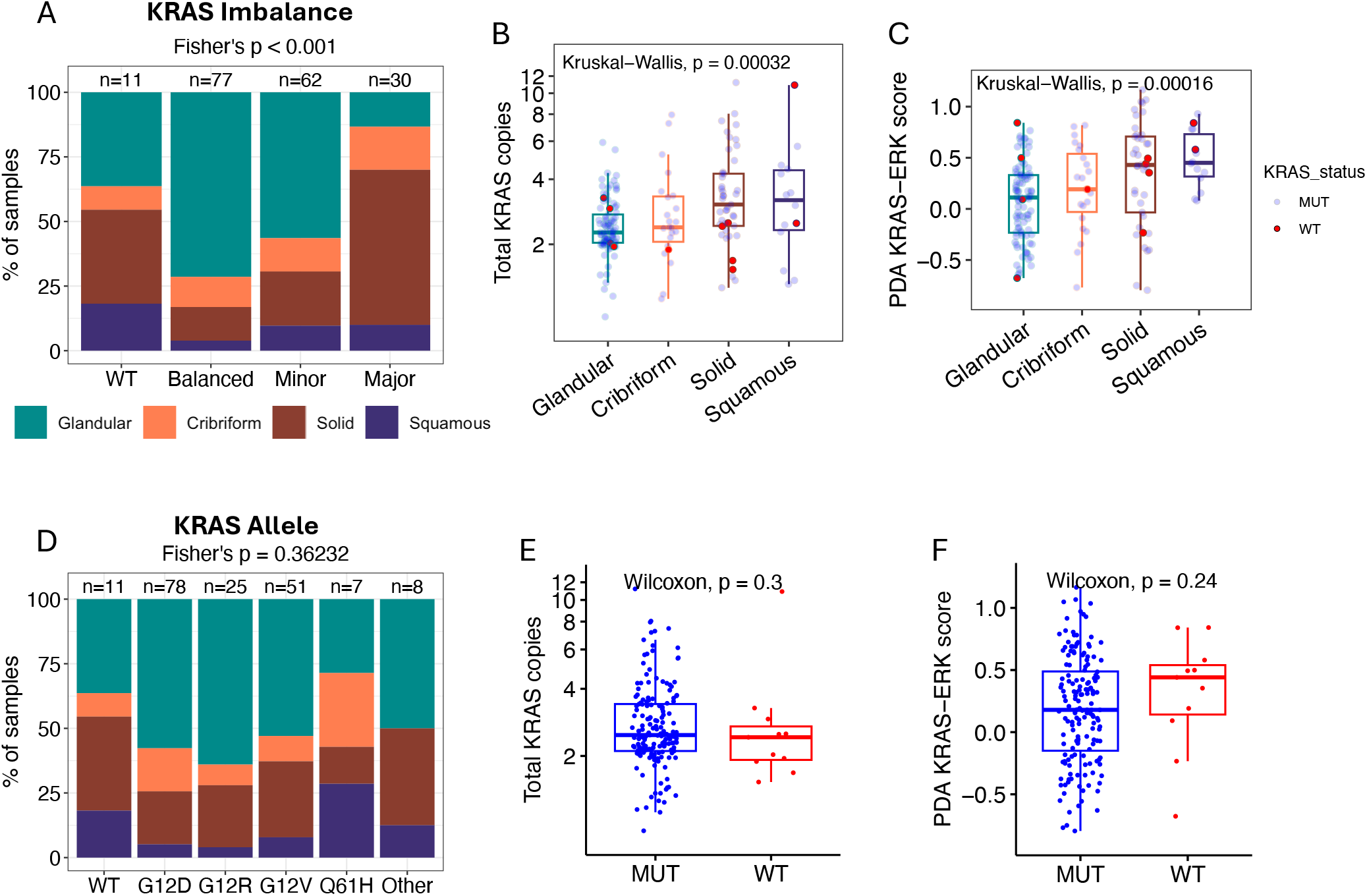
Relationship between KRAS status and morphological subtypes in advanced pancreatic ductal adenocarcinoma (PDAC). (A) Stacked bar plots showing the distribution of KRAS allelic imbalance across morphological subtypes, (B) Box plots depicting total KRAS copy number and, (C) the Gene Set Variation Analysis (GSVA) scores for the KRAS-ERK signalling pathway developed for PDAC by Klomp et al.^21^ (D) Stacked bar plot illustrating the distribution of KRAS allelic states by morphological subtype. (E,F) Box plots comparing KRAS total copy number and KRAS-ERK score (F) between KRAS mutant (MUT) and wild type (WT). All analyses were performed in advanced-stage cases.

### Clinical Associations between Morphological Classes in Resectable and Advanced Pancreatic Cancer

Morphological classes were strongly prognostic after surgical resection, with glandular tumours exhibiting significantly longer overall survival (OS) (p = 0.0005; **Figure 6A**). In the metastatic cohort, a trend toward shorter OS was observed for non-glandular compared with glandular tumours (p = 0.052; **Figure 6B**). If patients were divided using the 4 morphological classes, we observed no differences (**Supplementary Figure 3**)

**Figure 6.**
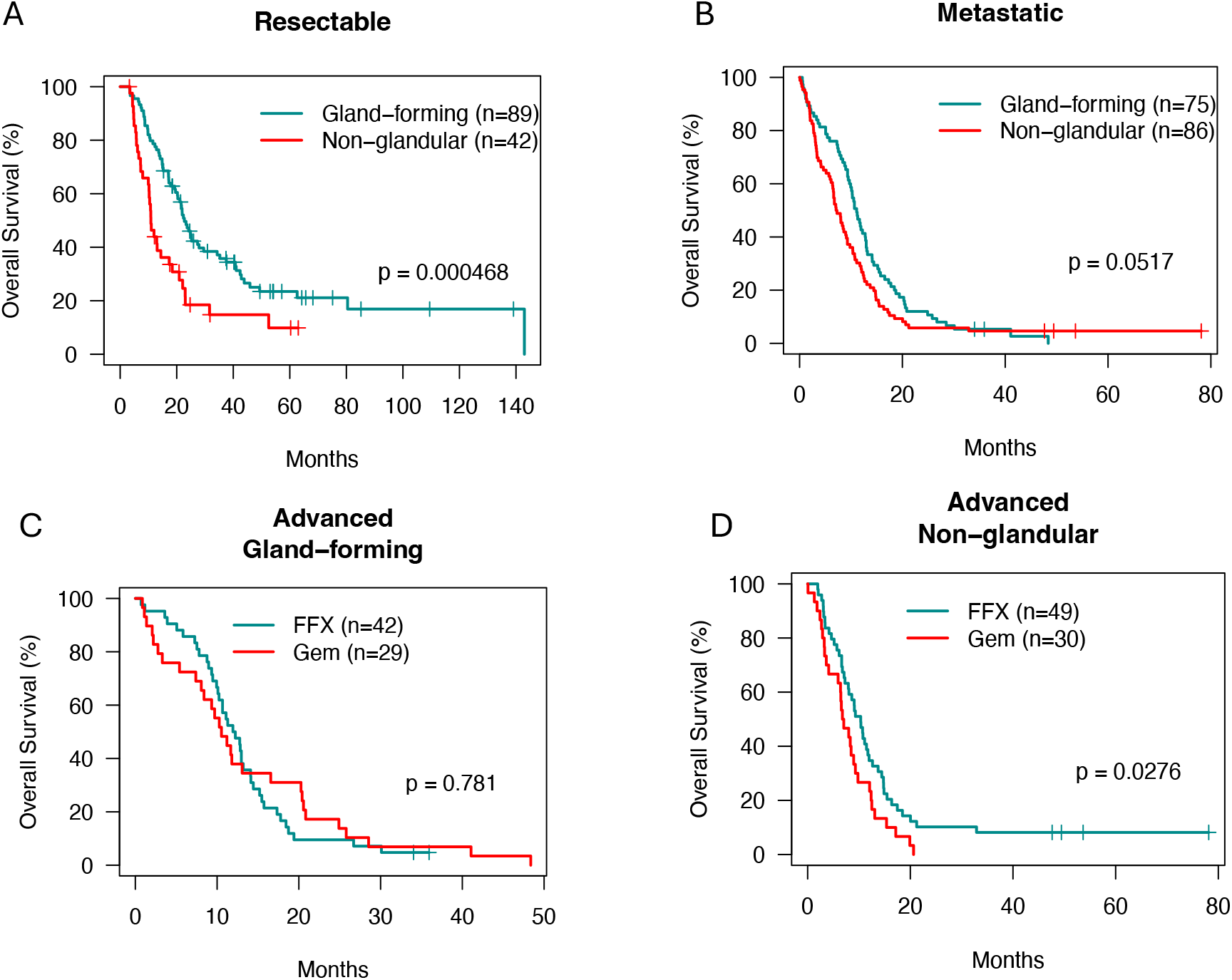
Survival associations of morphological subtypes and treatment response in pancreatic ductal adenocarcinoma (PDAC). (A, B) Kaplan–Meier survival curves comparing gland-forming and non-glandular tumours in patients with (A) resectable and (B) metastatic disease. (C, D) Kaplan–Meier survival curves comparing outcomes following FOLFIRINOX (FFX) and gemcitabine (Gem) treatment in patients with advanced disease, stratified by (C) gland-forming and (D) non-glandular tumours.

We did not find significant differences among morphological classes or detect a differential treatment benefit by chemotherapy regimen across survival or response, although these analyses were limited by small numbers (**Supplementary Figures 3B-G**). Together, these findings support that non-glandular morphologies are a hallmark of metastatic dissemination in PDAC.

## Discussion

Histological evaluation is central to the diagnosis and prognostication of PDAC, yet the molecular basis of its morphological heterogeneity remains poorly defined, particularly in advanced disease. In this study, we consolidated previously described morphological features into four classes, glandular, cribriform, solid, and squamous, and applied them to a large cohort including all PDAC stages with genomic and transcriptomic profiling. In contrast to prior studies focused predominantly on resected primary tumours or on transcriptional phenotypes independent of histology, our work systematically interrogates morphological classes across disease stages, integrates tumour-enriched genomic and transcriptomic data, and directly links histomorphology to molecular programs in advanced and metastatic PDAC.

Non-glandular morphologies were markedly enriched in metastatic lesions compared with primary pancreatic tumours, identifying them as a hallmark of metastatic PDAC. While primary tumours were predominantly glandular, metastatic biopsies showed a substantially higher proportion of cribriform, solid, and squamous architectures. Metastatic lesions also exhibited greater intra-tumour morphological heterogeneity, with mixed histological patterns observed more frequently than in primary tumours.

Integrating transcriptomic and histomorphological data, we identified distinctive basal-like states that are consistently restricted to non-glandular morphologies. Although these states express comparable levels of core basal signatures; they are defined by divergent biological programs including classical, pEMT, cell cycle, neural-like, keratinization, EMT and KRAS signaling. The unique transcriptional architecture of these subtypes supports their classification as independent entities, which may influence clinical outcomes and selection of therapeutic strategies.

Elevated KRAS signaling in solid and squamous tumours is linked to *KRAS* copy number gains and allelic imbalances, reinforcing the link between genomic instability and oncogenic drivers^13,22^. Our findings further show that solid morphologies exhibit polyploidy and high KRAS signaling, while yielding the highest EMT scores. This further validates the association between EMT and genomic instability in PDAC recently established by Perelli *et al*.^23^. Ultimately, these results highlight the interplay between KRAS signaling, genomic instability, EMT and morphological progression. Beyond *KRAS*, non-glandular progression was associated with *MYC* copy number gains, which agrees with previous correlations with metastatic progression^8,13^ and squamous differentiation^3^. We did not find other major genomic differences between the morphological classes. This observation is supported by the study of Hayashi *et al*. on squamous morphology, where they observed that metastases and primary tumours from the same patients may present divergent morphologies, despite similar genomes^3^. Together, these findings suggest that morphological diversity in PDAC is primarily driven by transcriptional reprogramming rather than widespread differences in mutational landscapes.

One limitation of this study is the cross-sectional nature of the sampling, which precludes direct inference about the temporal relationship between morphological states and metastatic dissemination. Our analysis captures associations between morphology and disease context rather than the evolutionary trajectory of individual tumours. In addition, metastatic samples were primarily obtained from biopsies, which may incompletely capture spatial heterogeneity within lesions. Future studies integrating longitudinal sampling with matched primary and metastatic lesions will be needed to define the origins and stability of morphological states in PDAC.

Together, these results show that morphological classes correspond to distinct biological states in PDAC. By integrating histology with transcriptomic and genomic profiling across primary and metastatic tumours, we identify molecular programs associated with glandular, cribriform, solid, and squamous morphologies. Non-glandular morphologies are enriched in metastatic disease and associate with genomic instability and increased KRAS–ERK signalling. These findings link tumour morphology with metastatic dissemination and distinct transcriptional and genomic features in PDAC. Our observations regarding the association between morphological classes and KRAS doses support further investigation of tumour morphology as a hypothesis-generating biomarker for KRAS-targeted therapies.

## Data availability

The WGS of germline DNA and tumour tissues and RNA-seq of tumour tissues can be found in the European Genome-phenome Archive (EGA): EGAD00001004551, EGAD00001006262, EGAD00001007571, EGAD00001006261, EGAD00001009409.

## Contributions

Conceptualized the project (EFF,YF,FN,RG), Collected/organized patient samples/clinical data (SR, DB, SH,RD, ET, SG, JK), processing samples (KN, S-BL, KN), managed and analyze raw sequencing data (GJ, MCH-S-Y, AZ), performed most of the analysis and create figures (YF, EFF), provided additional biostatistics analysis (TO), Annotated all histological slides (AE, MM), wrote the manuscript (EFF, YF, FN, RG), Provided feedback (KN, SF, JW, AD).

## Competing interests

The authors declare no competing interests.

## Acknowledgements and Funding

This study was conducted with the support of the Ontario Institute for Cancer Research (OICR) through funding provided by the Government of Ontario, the Wallace McCain Centre for Pancreatic Cancer supported by the Princess Margaret Cancer Foundation, the Terry Fox Research Institute Marathon of Hope Cancer Centres Network, the Canadian Cancer Society Research Institute, Pancreatic Cancer Canada and the Canadian Cancer Society Breakthrough Team Grant (CCS grant #707708). The study was also supported by a charitable donation from the Canadian Friends of the Hebrew University (Alex U. Soyka). We acknowledge the contributions of the UHN Cancer Biobank and members of the OICR Diagnostic Development (Tissue Portal; oicr.on.ca/programs/diagnostic-development/), Genomics (genomics.oicr.on.ca) and PanCuRx programs for sample management, sequencing and data analysis. Figure 1A was created in BioRender.

## Methods

### Pancreatic cancer patients

Written informed consent was obtained from all patients (n = 348). Basic demographic data is described in Supplementary Table 1. Most of the patient samples were collected at Princess Margaret Cancer Centre at the University Health Network (Toronto, Canada). Tumours from resectable cases were obtained from the University Health Network (UHN) Biospecimens Program, and the ones from advanced cases were obtained from the COMPASS trial (no. NCT02750657). As part of previous studies^8,14,25,26^, some tumours were obtained from Sunnybrook Health Sciences Centre (Toronto), Kingston General Hospital (Kingston), McGill University (Montreal), Mayo Clinic (Rochester), Massachusetts General Hospital (Boston) and Sheba Medical Centre (Tel Aviv). Approval for the study was obtained through the UHN Research Ethics Board (nos. 15-9596, 13-6377, 18-5116, 20-5594, 21-5648 and 32517).

### Tumour Whole-Genome Sequencing and Analysis

For tumour enrichment, laser capture microdissection (LCM) was performed on frozen biospecimens. Genomic DNA was extracted from the microdissected tumours using the Gentra Puregene Tissue kit components (Qiagen) and was quantified using the Qubit dsDNA High Sensitivity Kit (Invitrogen). For the library preparation of whole-genome Sequencing (WGS), either the NEBNext DNA Sample Prep Master Mix Set (New England Biolabs), the Nextera DNA Sample Prep Kit (Illumina) or the KAPA Library Preparation Kits (Roche) was used. Established libraries were quantified with the Illumina Eco Real-Time PCR Instrument using KAPA Illumina Library Quantification Kits (Roche). Paired-end WGS was conducted with the Illumina HiSeq 2000/2500 platform or the NovaSeq 6000 with a target coverage of 50x. For paired normal reference genomic DNA, usually from buffy coat or normal tissues, a similar protocol was used for DNA extraction and library preparation, and WGS was conducted with a targeted coverage at 35x.

Raw sequencing reads were aligned to the human reference genome (hg38) using Burrow-Wheeler Aligner (BWA, version 0.7.17)^27^. Germline variant calling was performed using Genome Analysis Tool Kit (GATK, version 4.1.2)^28^. Somatic single nucleotide variants (SNVs) were identified as the intersection of calls by Strelka2 (version 2.9.10)^29^ and MuTect2 (version 4.1.2)^30^. Indels were identified using the consensus between two of four callers: Strelka2, MuTect2, SVaBA (v134)^31^ and DELLY2 (version 0.8.1)^32^. Somatic structural variants were identified using the consensus of 2 of 3 callers: SVaBA, DELLY2, and Manta (version 1.6.0)^33^. HMMcopy (version 0.1.1) (https://github.com/shahcompbio/hmmcopy_utils) and CELLULOID^14^ were used to call copy number segments, tumour cellularity, and ploidy.

### Genomic features

Patients’ tumour samples were classified as homologous recombination deficient (HRD) based on established genomic hallmarks, as previously described^34,35^. These hallmarks comprised alterations in HRD-associated genes, single-nucleotide variant (SNV) and structural variant (SV) burdens, frequencies of ≥4 bp deletions, 100 bp–10 kb deletions, and 10 kb–1 Mb duplications, as well as the C>T substitution ratio, ≥4 bp deletion ratio, and 100 bp–10 kb deletion ratio. Tumours exhibiting six or more HRD hallmarks were classified as HRD, while those with three to five hallmarks underwent manual curation.

*KRAS* subtypes were defined according to previously published criteria^20^. Tumours lacking nonsynonymous SNVs in *KRAS* were classified as wild type (WT). In *KRAS*-mutant tumours, the ratio of mutant to wild-type copy number at the *KRAS* locus was calculated. Tumours were categorised as balanced (ratio = 1), minor imbalanced (ratio >1 and <3), or major imbalanced (ratio ≥3).

### Tumour RNA Sequencing and Analysis

RNA was extracted from the microdissected tumours using the PicoPure RNA Isolation Kit (ThermoFisher Scientific) and quantified using the Qubit dsRNA High Sensitivity Kit (Invitrogen). Quality control was conducted on extracted RNA using the RNA ScreenTape Assay on the 2200 TapeStation Nucleic Acid System (Agilent Technologies). RNA with RNA integrity number > 7 was used for library preparation using the TruSeq RNA Access Library Sample prep kit (Illumina) or the TruSeq Stranded Total RNA Library Prep Gold kit (Illumina). Established libraries were quantified using the KAPA Illumina Library Quantification Kit (Roche). Paired-end RNA sequencing was carried out on the Illumina HiSeq 2500 platform or the NovaSeq 6000 with a target of a minimum of 30 million unique mapped reads.

Reads were aligned to the human reference genome (hg38) and transcriptome (Ensembl v.100) using STAR v.2.7.4a^36^. Gene raw counts were obtained by HTSeq (version 2.0.4)^37^, and were used for differential expression analysis with DESeq2 (version 1.30.1)^38^, where the log2FC shrinkage with type = ‘ashr’ was used for effect size shrinkage^39^. To identify the signature genes for the morphological subtypes, a One vs. Rest (OVR) approach was used, where tumours with each morphology were compared to the rest to obtain the genes that were up-regulated in this morphology. This process was repeated for all 4 morphological subtypes. Up-regulated genes that were unique to one morphology (i.e., did not occur in other morphologies) were defined as the signature genes for that morphology. Overrepresentation analysis with GO terms^40^ was conducted on the signature genes to identify enriched gene sets in each morphology using clusterProfiler (version 3.18.1)^41^. Gene lists from PDAC RNA signatures were extracted from publications^6,10– 13,42,43^, and these signatures were quantified by gene set variation analysis (GSVA)^44^.

### Histopathological review

All the digitally scanned Haematoxylin and eosin (H&E) stained slides from all patients with PDAC were first reviewed by a pathologist (MM) blinded to clinical, original pathology report and RNA-Seq data. The tissue origin (biopsy site), tumour type and subtype based on the World Health Organization (WHO) tumour classification was scored^9^. All digitally scanned hematoxylin and eosin (H&E)–stained slides from patients diagnosed with PDAC were independently reviewed using QuPath software, by two board-certified pathologists (MM and AE), who were blinded to clinical data, original pathology reports, and RNA sequencing results. For each case, the tissue origin (biopsy site) and tumour type and subtype were determined according to the criteria outlined in the WHO Classification of Tumours. Tumour cellularity, necrosis, and stromal content were assessed semi-quantitatively and recorded as percentages of the total tumour area. In addition, tumours were evaluated for previously described morphologic subtypes as reported by Kalimuthu *et al*.^2^ and Di Chiaro *et al*.^4^. Tumours were further classified according to our predefined morphologic categories: glandular, cribriform, solid, and squamous. For each slide, the relative proportion of each morphologic pattern was estimated as a percentage of the total tumour area. The dominant morphologic pattern was defined as the subtype comprising the largest percentage

## Supplementary Figure Legends

**Supplementary Figure 1.**
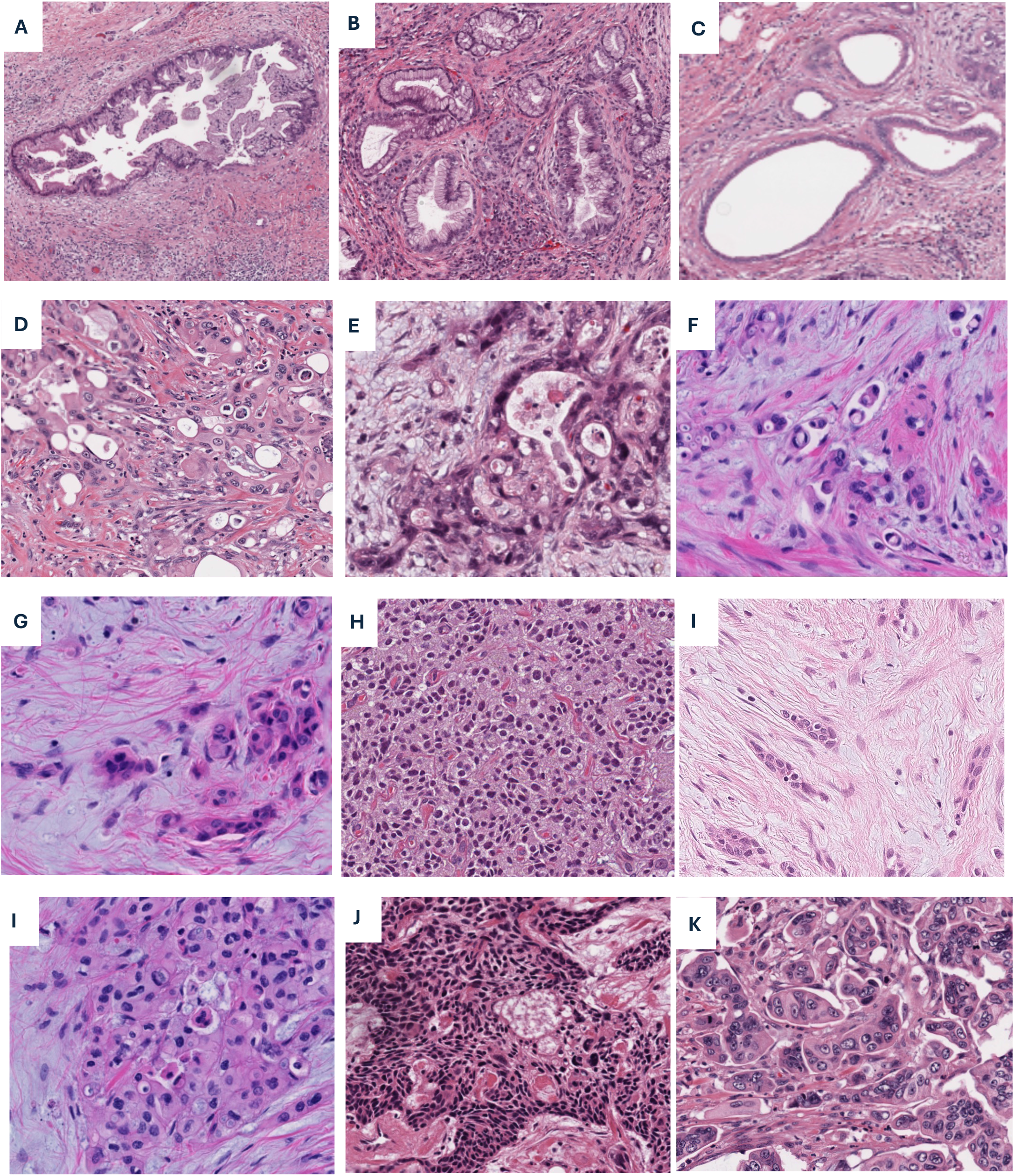
Morphological subtypes of pancreatic ductal adenocarcinoma in the H&E stain. (A-C) Glandular class. Tumours characterized by well-formed glandular architecture with preserved luminal differentiation and epithelial polarity. (A) Large, well-formed patent gland with intraluminal papillary projections, lined by well-differentiated foveolar-type epithelium. (B) Smaller infiltrative glands lined by well-differentiated foveolar-type epithelium. (C) Variably sized glands lined by moderately differentiated pancreaticobiliary-type epithelium. (D-F) Cribriform class. Tumours lacking well-formed glandular architecture but retaining glandular cytomorphology, demonstrating progressive architectural disorganization. (D) Cohesive clusters of moderately differentiated tumour cells with punched-out spaces secondary to central necrosis. (E) Poorly formed, angulated gland-like structures with minimal to absent intervening stroma. (F) Predominantly discohesive single tumour cells with signet ring morphology. (G-I) Solid class. Tumours showing complete loss of glandular architecture and glandular cytomorphology. (G) Small clusters of tumour cells lacking glandular differentiation. (H) Confluent sheets of poorly differentiated tumour cells. (I) Tumour cells arranged in cords, reflecting a non–gland-forming growth pattern. (J-L) Squamous class. Tumours exhibiting squamous differentiation with variable degrees of maturation. (J) Clusters of polygonal cells with abundant eosinophilic cytoplasm, distinct intercellular bridges, and scattered dyskeratotic cells. (K) Sheets of smaller tumour cells with dense eosinophilic cytoplasm and minimal keratinization. (L) Small clusters of variably sized tumour cells with eosinophilic cytoplasm, enlarged nuclei, and prominent nucleoli.

**Supplementary Figure 2.**
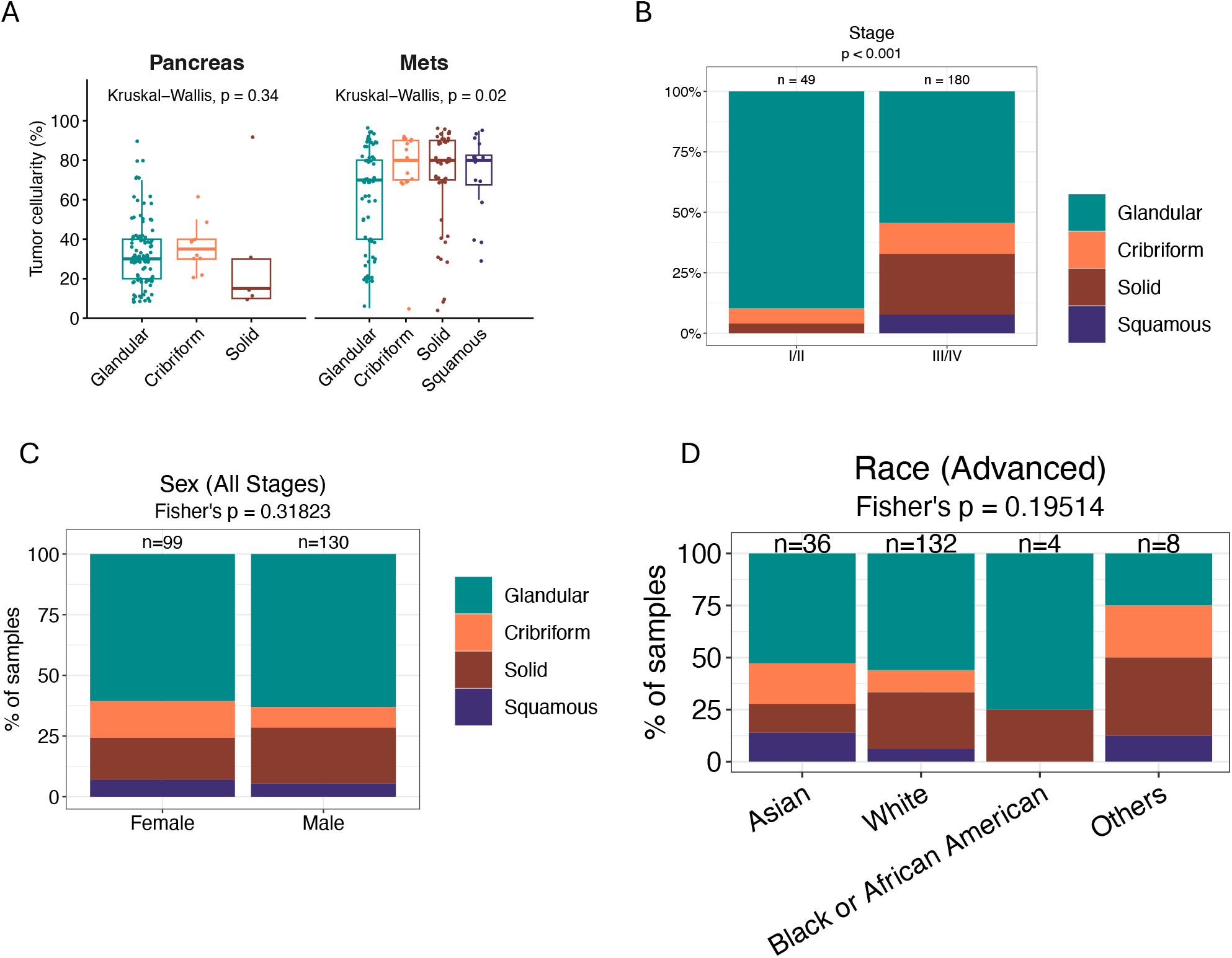
(A) Tumour cellularity (prior to laser micro-dissection) between primary pancreatic and metastatic tumours across morphological subtypes. Stacked bar plots indicating the percentage of samples for each morphological subtype, categorized by stage (B), sex (C) and race (D).

**Supplementary Figure 3.**
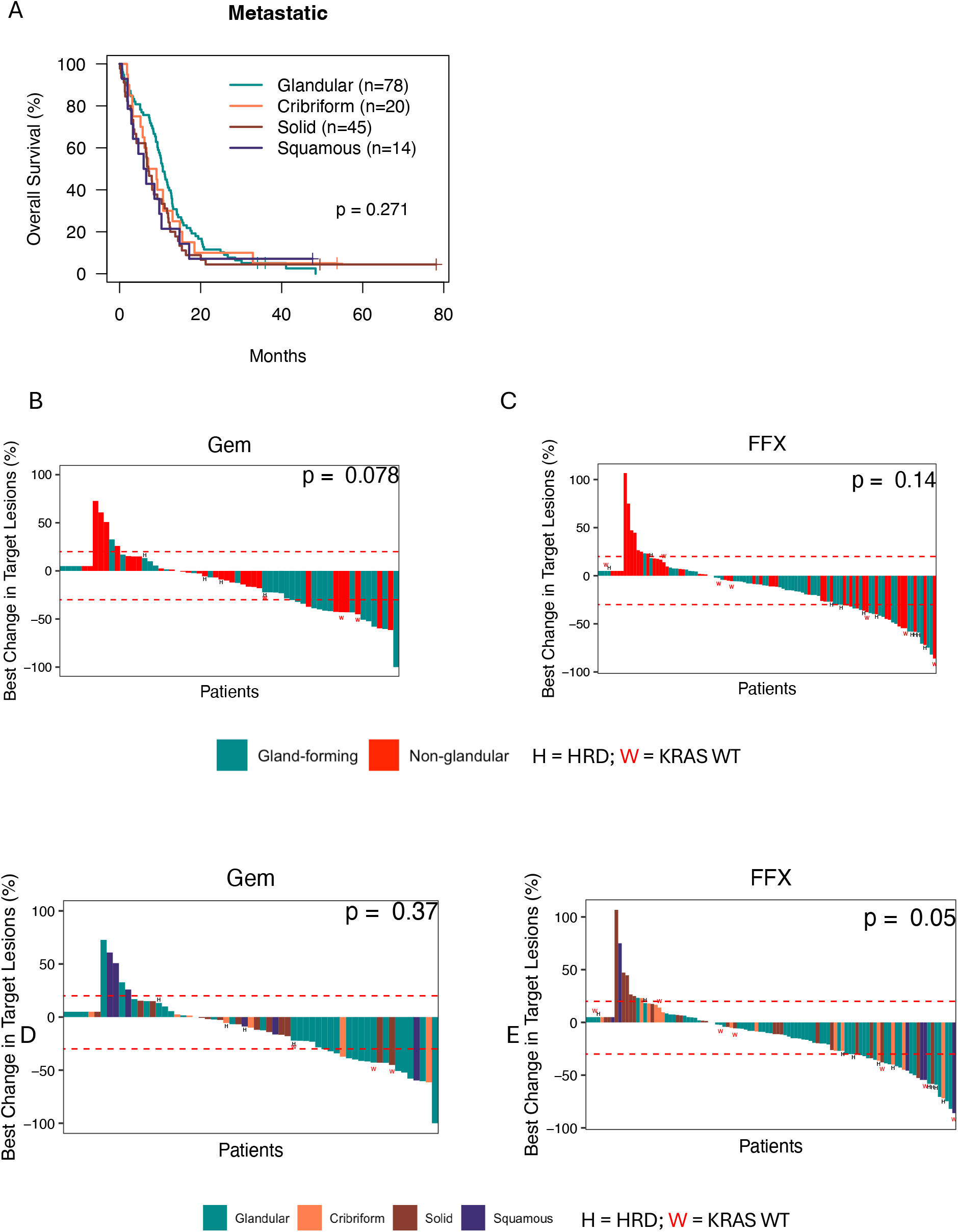
(A) Kaplan–Meier survival curves comparing glandular, cribriform, solid, and squamous tumours in patients with metastatic disease. (B–E) Waterfall plots showing responses of glandular and non-glandular tumours to first-line gemcitabine (B) or FOLFIRINOX (C), and responses of glandular, cribriform, solid, and squamous tumours to first-line gemcitabine (D) or FOLFIRINOX (E), according to RECIST criteria, in advanced cases (locally advanced and metastatic). Dashed red lines indicate thresholds for partial response (PR, −30%) and progressive disease (PD, +20%). Each bar represents an individual patient and is coloured by morphology. Patients without evaluable tumour size measurements are shown to the left (not evaluable, NE). Homologous recombination deficiency (H) and KRAS wild-type (W) cases are labelled. The Wilcoxon signed-rank test was used to compare changes in tumour size between gland-forming and non-glandular tumours.

**Supplementary Figure 4.**
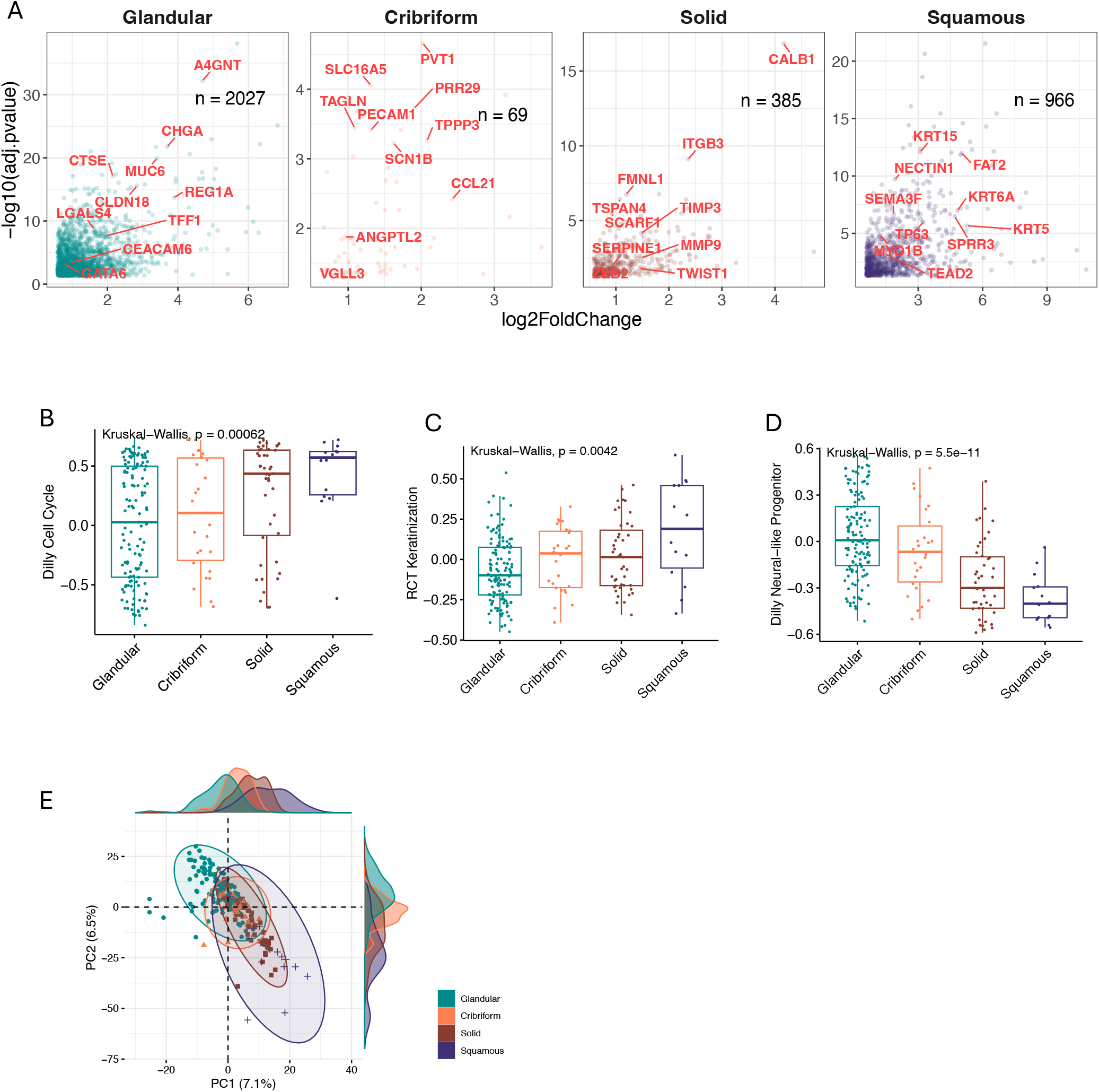
(A) Morphology-specific genes identified using a one-versus-rest differential expression analysis (Methods). Fold change indicates the expression level of each gene in the indicated morphological class relative to all other tumours. Gene Set Variation Analysis (GSVA) scores for (B) cell cycle from Dilly *et al*.^10^, (C) keratinization from Reactome^24^ and (D) neuronal-like progenitor from Dilly *et al*.^10^. (E) Principal component analysis (PCA) of morphology-specific genes illustrating a transcriptional trajectory from glandular (green) to cribriform (orange), solid (brown), and squamous (purple) tumours.

## Notes

### Competing Interest Statement

The authors have declared no competing interest.

